# Maternal SMCHD1 controls both imprinted *Xist* expression and imprinted X chromosome inactivation

**DOI:** 10.1101/2021.10.21.465360

**Authors:** Iromi Wanigasuriya, Sarah A Kinkel, Tamara Beck, Ellise A Roper, Kelsey Breslin, Heather J Lee, Andrew Keniry, Matthew E Ritchie, Marnie E Blewitt, Quentin Gouil

## Abstract

Embryonic development is dependent on the maternal supply of proteins through the oocyte, including factors setting up the adequate epigenetic patterning of the zygotic genome. We previously reported that one such factor is the epigenetic repressor SMCHD1, whose maternal supply controls autosomal imprinted expression in mouse preimplantation embryos and mid-gestation placenta. In mouse preimplantation embryos, X chromosome inactivation is also an imprinted process. Combining genomics and imaging, we show that maternal SMCHD1 is required not only for the imprinted expression of *Xist* in preimplantation embryos, but also for the efficient silencing of the inactive X in both the preimplantation embryo and mid-gestation placenta. These results expand the role of SMCHD1 in enforcing the silencing of Polycomb targets. The inability of zygotic SMCHD1 to fully restore imprinted X inactivation further points to maternal SMCHD1’s role in setting up the appropriate chromatin environment during preimplantation development, a critical window of epigenetic remodelling.

## Introduction

X chromosome inactivation (XCI) in female mammals is a paradigm of epigenetic regulation, where hundreds of genes on a single chromosome coordinately undergo silencing (***Lyon, 1961, 1962***). In the common ancestor of therian mammals, XCI evolved as a mechanism of sex chromosome dosage compensation, balancing female X-linked expression at levels similar to males possessing only one X chromosome (***Lyon, 1963; Reik and Lewis, 2005***). The ancestral form of XCI is potentially imprinted, with preferential silencing of the paternal X, whereas random X inactivation is proposed to be derived (***Reik and Lewis, 2005; Deakin et al., 2009***). In marsupials, imprinted X inactivation is maintained in all tissues (***Cooper et al., 1971***) whereas in rodents or cattle it only persists in extraembryonic tissues that gives rise to the placenta, while the embryo-proper reactivates the paternal X before random inactivation of either the maternal or paternal chromosome takes place (***Okamoto et al., 2011***). In humans, only random X inactivation occurs.

In mice, imprinted X inactivation originates in the preimplantation embryo (***Okamoto and Heard, 2006***). Systematic silencing of the paternal X is caused by a Polycomb-mediated repressive imprint laid down during oogenesis, which prevents the long non-coding RNA *Xist* from being expressed (***Tada et al., 2000; Chiba et al., 2008; Inoue et al., 2017***). Paternal expression of *Xist* thus leads to silencing of the paternal X (***Heard et al., 2004***). Maternal effect genes that control the epigenetic patterning of the oocyte and early zygote are important for the correct imprinted expression of *Xist* (***Inoue et al., 2017; Harris et al., 2019; Mei et al., 2021; Chen et al., 2021***). The genomic region surrounding *Xist* houses multiple positive and negative regulators of *Xist* expression, and is termed the X-inactivation centre (***Galupa and Heard, 2018***).

We previously established that the maternal supply of Structural Maintenance of Chromosome Hinge Domain containing 1 (SMCHD1) regulates some of the Polycomb-dependent imprinted genes on autosomes (***Wanigasuriya et al., 2020***). Both *Xist* and autosomal Polycomb-dependent imprinted genes are non-canonical imprinted genes, as they rely on Polycomb marks for their imprinted expression rather than DNA methylation as canonical imprinted genes do. Based on this role of maternal SMCHD1 and the known involvement of zygotic SMCHD1 in XCI (***Blewitt et al., 2008; Gendrel et al., 2012***), we investigated whether maternal SMCHD1 also played a role in regulating the imprinted expression of *Xist*, and whether it affected silencing of the inactive X. Through epigenomic and imaging analyses of preimplantation embryos and mid-gestation placentae, we show that SMCHD1 is also a maternal effect gene with regard to imprinted X chromosome inactivation.

## Results

### Maternal deletion of *Smchd1* results in aberrant *Xist* expression from the maternal allele

To determine the role of maternal SMCHD1 on imprinted *Xist* expression and X inactivation, we ablated *Smchd1* in mouse oocytes using the MMTV-Cre transgene and crossed the dams with wild-type sires from a different strain to allow allele-specific analyses (Figure 1a, as reported in (***Wanigasuriya et al., 2020***)). We analysed single-embryo methylome and transcriptome data for *Smchd1*^*mat*Δ^ and control *Smchd1*^*wt*^ E2.75 embryos (16–32 cells), when zygotic SMCHD1 only just starts to accumulate (***Wanigasuriya et al., 2020***).

**Figure 1.**
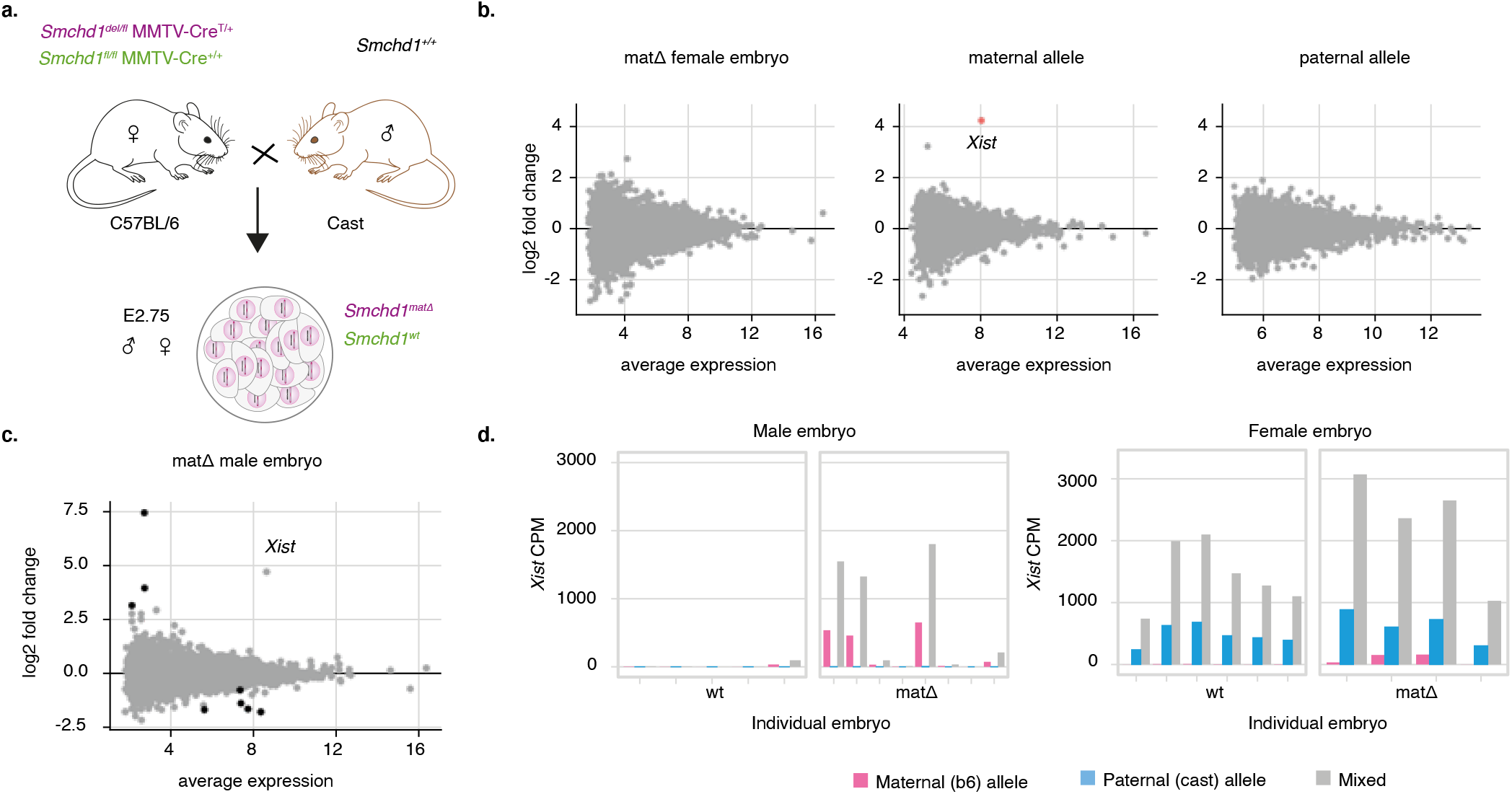
Maternal deletion of *Smchd1* results in aberrant *Xist* expression from the maternal allele in both male and female E2.75 preimplantation embryos. (a) Schematic of genetic crosses for maternal deletion of *Smchd1*. (b) Genome-wide differential expression in *Smchd1*^*mat*Δ^ female embryos vs wt, before haplotyping and after separating maternal and paternal alleles. Average expression is in log2 counts per million (cpm). Only maternal *Xist* is significantly differentially expressed (adjusted p-value = 6e-4). (c) Genome-wide differential expression in *Smchd1*^*mat*Δ^ male embryos vs wt, without haplotyping. Significant genes are coloured black (5% FDR). *Xist* is not significant (adjusted p-value = 0.17) due to partial penetrance in *Smchd1*^*mat*Δ^ samples, but has a large log_2_ fold change (4.74). (d) *Xist* expression in individual male and female wt and *Smchd1*^*mat*Δ^ E2.75 embryos. CPM: counts per million (of total library size before haplotyping). “Mixed” counts refer to counts without haplotyping. Females: n = 6 wt and 4 mat.Δ; males: n = 5 wt and 8 mat.Δ. **Figure 1–Figure supplement 1.** *Rhox9* expression in individual male and female wt and *Smchd1*^*mat*Δ^ E2.75 embryos. cpm: counts per million (of total library size before haplotyping). “Mixed” counts refer to counts without haplotyping. Females: n = 6 wt and 4 mat.Δ; males: n = 5 wt and 8 mat.Δ. **Figure 1–Figure supplement 2.** DNA methylation at the X inactivation center in female and male E2.75 wt embryos and *Smchd1*^*mat*Δ^ embryos. Histogram tracks show methylation in single embryos at individual CpG sites (0–100%, 4 wild-type and 4 mat.Δembryos shown for each sex). Note that coverage in single-embryo whole-genome bisulfite sequencing is sparse, only 0–2X. The aggregate line plots show the average methylation per genotype across 10 kb windows (sliding by 5 kb, 0–100%). Females: n = 6 wt and 4 mat.Δ; males: n = 5 wt and 8 mat.Δ.

We previously reported very little genome-wide differential expression in male preimplantation embryos without maternal SMCHD1 (***Wanigasuriya et al., 2020***). Consistent with that, there was also very little genome-wide differential expression in female *Smchd1*^*mat*Δ^ embryos compared to wild-type controls (Figure 1b and c). Without haplotyping RNA-seq counts, there were no significantly differentially expressed genes at the 5% FDR threshold in female *Smchd1*^*mat*Δ^ embryos, and only 8 in male embryos (upregulated *Rhox9, E330020D12Rik* and *Cdc42bpa*, and downregulated *Hspa5, Akr1a1, Hsp90b1, Calr* and *Pdia6*, Supplementary Tables 1-2). *Rhox9* had the strongest log-fold change (7.5). It is an imprinted gene (***MacLean et al., 2011***), subject to H3K27me3- and DNA methylation-mediated repression (***Berletch et al., 2013***), and part of a clustered gene family: recurrent characteristics among SMCHD1 targets. In females *Rhox9* was filtered out of the differential expression analysis because of low counts, but upon manual investigation the data also supported maternal *Rhox9* upregulation in *Smchd1*^*mat*Δ^ samples (Figure 1-Figure supplement 1).

Both in the male total expression and female allele-specific expression, the maternal copy of *Xist* was a striking outlier with a high level of expression and large upregulation in *Smchd1*^*mat*Δ^ samples (log_2_ fold changes of 4–5). Maternal *Xist* is normally silenced in the early embryo due to a Polycomb-mediated imprint (***Inoue et al., 2017; Harris et al., 2019; Mei et al., 2021; Chen et al., 2021***). Here maternal *Xist* was activated both in the female and male *Smchd1*^*mat*Δ^ embryos (Figure 1b and c). Although striking, maternal *Xist* loss of imprinting was not completely penetrant: 4 out of 8 male and 3 out of 4 female *Smchd1*^*mat*Δ^ embryos showed increased levels of maternal *Xist* expression (Figure 1d). This high variability in expression explained why *Xist* was not detected as statistically significant in the male embryo differential expression analysis (44th ranked gene, FDR=0.17). In the female embryos, paternal *Xist* expression still outweighed expression from the maternal allele. This could be due to low levels of zygotic SMCHD1 beginning to accumulate around E2.75 (***Wanigasuriya et al., 2020***) and partial silencing of maternal *Xist*, or additional repression mechanisms independent of SMCHD1. The partial penetrance could similarly be due to zygotic SMCHD1 expression, or it may reflect true biological variation in the response to maternal SMCHD1 ablation.

At the blastocyst stage the X-inactivation centre is partially methylated (***Prissette, 2001; McGraw et al., 2013***), so we asked whether the failure to silence maternal *Xist* could be due to a failure to acquire DNA methylation at the X-inactivation centre, using our E2.75 embryo DNA methylation data. However, the whole X-inactivation centre including the maternal *Xist* promoter and Xite/DXPas34 remained unmethylated in male and female wild-type E2.75 embryos, and there was no difference in the *Smchd1*^*mat*Δ^ embryos that displayed loss of imprinting (Figure 1 supplement 1). The loss of *Xist* silencing was therefore not linked to a defect in DNA methylation.

### Imprinted X chromosome inactivation is altered in *Smchd1*^*mat*Δ^ morulae

We then asked whether maternal *Xist* expression was functionally linked to the silencing of the maternal X chromosome. Although at the genome-wide level few individual genes passed the significance threshold for differential expression (Figure 1 b and c), the distribution of log_2_ fold changes (matΔvs wt) shifted significantly for X-linked genes (Figure 2a). In *Smchd1*^*mat*Δ^ males, the genes from the maternal X chromosome tended to be downregulated (mean log_2_ fold change = -0.16, equivalent to a reduction by 11%, p-value = 2.4e-5), consistent with aberrant *Xist*-mediated silencing. By contrast in *Smchd1*^*mat*Δ^ females, the alleles on the maternal X were not significantly downregulated (mean log_2_ fold change = -0.094, p-value = 0.18), meaning that the partial maternal *Xist* loss of imprinting did not trigger detectable silencing of the maternal X. This was surprising, however we cannot rule out that some Xm silencing is occurring but the effect is too small to detect with our current power. On the other hand, the paternal X appeared slightly upregulated compared to the wild-types (mean log_2_ fold change = 0.20, equivalent to an increase by 15%, p-value = 7.6e-6). When subsetting the embryos that specifically showed *Xist* loss of imprinting, the effects were stronger for the males (X downregulation by 25% on average) and unchanged for the females (Figure 2 supplement 1). We next asked whether the compromised paternal X silencing observed in *Smchd1*^*mat*Δ^ female embryos corresponded to a complete or partial loss of X chromosome inactivation. At E2.75, imprinted paternal X inactivation is normally ongoing and does not yet affect all the genes on the paternal X chromosome (***Patrat et al., 2009; Borensztein et al., 2017***). Accordingly, in our data the paternal X in the wild-type females was downregulated by 45% on average (Figure 2b). Therefore, the paternal X repression observed in *Smchd1*^*mat*Δ^ was only partially compromised compared to the wild-type scenario. Taken together, these results indicate that: 1) aberrant maternal *Xist* expression can lead to partial Xm silencing, 2) initiation of X chromosome inactivation can occur in the absence of maternal SMCHD1 (Xm silencing in males, remaining Xp silencing in females). Finally, incomplete Xp silencing compared to the wild-type scenario may have multiple alternative explanations: perhaps maternal SMCHD1 still contributes to part of these early stages of imprinted X inactivation; biallelic Xist expression could titer the silencing machineries between the two X chromosomes; or biallelic Xist expression might slow the rate of development and/or X chromosome inactivation.

**Figure 2.**
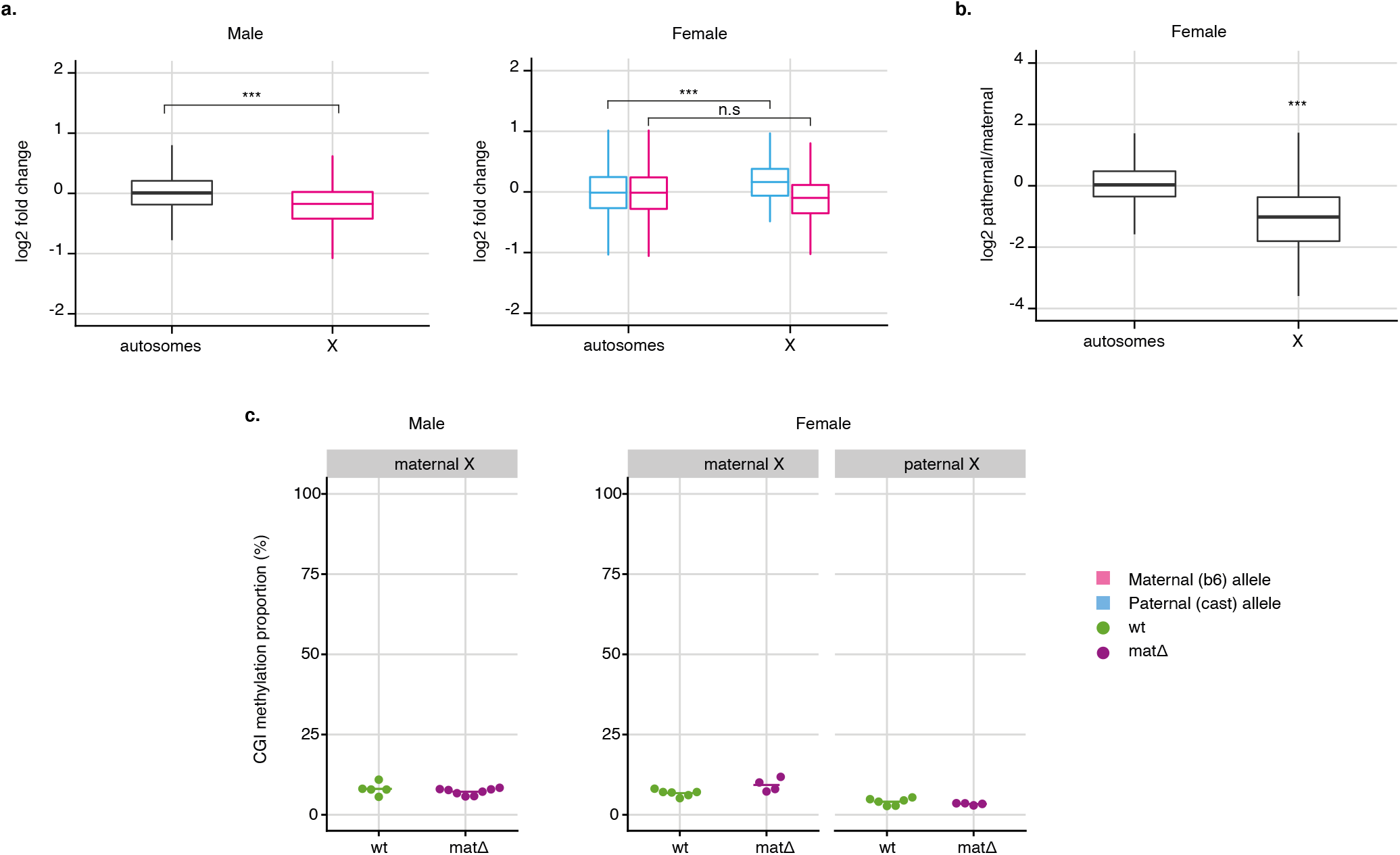
Maternal deletion of *Smchd1* results in aberrant XCI in male and perturbed imprinted XCI in female E2.75 preimplantation embryos. (a) Distribution of *Smchd1*^*mat*Δ^ vs wt gene expression log_2_ fold changes on autosomes and the X chromosome for male and female E2.75 embryos. For females, results of the allele-specific differential expression analysis are shown, with paternal alleles in blue and maternal alleles in pink. Two-sample t-tests; male: p = 2.4e-5; female paternal allele p = 7.6e-6; female maternal allele p = 0.18. (b) Distribution of paternal over maternal log2 expression ratios in wt female E2.75 embryos. Paternal X-linked genes are significantly repressed (p = 1.5e-4, one-sample t-test). (c) Average CpG island (CGI) methylation on the X chromosomes of individual *Smchd1*^*wt*^ and *Smchd1*^*mat*Δ^ male and female E2.75 embryos. Females: n = 6 wt and 4 matΔ; males: n = 5 wt and 8 matΔ. T-tests, males maternal X: p-value = 0.4; females maternal X: ; females maternal X: p-value = 0.1; females paternal X: p-value = 0.2. **Figure 2–Figure supplement 1.** Distribution of *Smchd1*^*mat*Δ^ vs wt gene expression log_2_ fold changes on autosomes and the X chromosome for male and female E2.75 embryos, retaining only matΔembryos with loss of *Xist* imprinting. **Figure 2–Figure supplement 2.** Whole-genome differential methylation analysis between female *Smchd1*^*mat*Δ^ and wild-type E2.75 embryos. For CpG islands (CGIs, 13k regions), promoters (−4 kb to +1 kb regions, 52k regions) and 10-kb windows (sliding by 5 kb, 500k regions), the average methylation level in wild types is plotted against the average methylation in *Smchd1*^*mat*Δ^ embryos. Significant Differentially Methylated Regions (DMRs, FDR < 5% and absolute difference in methylation > 20%) are coloured in red (hypermethylation) or blue (hypomethylation). Females: n = 6 wt and 4 matΔ.

To further investigate the stage and mechanisms of XCI in the E2.75 embryos, we analysed CpG island methylation on each allele of the X chromosome. In wild-type female embryos, average CGI methylation on the paternal inactive X remained low and similar to that of the maternal X and the male X (Figure 2c). CpG island methylation in *Smchd1*^*mat*Δ^ embryos was indistinguishable from the wild types. These results imply that maternal SMCHD1 has no role in the methylation of the inactive X at this time.

Genome-wide, there was little evidence of differential methylation between wild-type and *Smchd1*^*mat*Δ^ female E2.75 embryos (Figure 2 supplement 2), similar to what we reported in male embryos (***Wanigasuriya et al., 2020***). Methylation at CpG islands was very low, similar to the CGIs of X chromosomes (Figure 2c), and there were no significant differentially methylated CGIs (Figure 2 supplement 2). Across gene promoters, methylation levels were more broadly distributed but highly consistent between wild-type and *Smchd1*^*mat*Δ^ embryos. There was a relative excess of hyper-methylated promoters in the *Smchd1*^*mat*Δ^ samples, but with mild (<30%) differences and making up only 0.5% of all promoters (246 hypermethylated, 10 hypomethylated, out of 52k promoters) with no overlap with differentially methylated promoters in males. Over 10 kb windows sliding across the whole genome, 445 were hypermethylated including 42 also hypermethylated in the males, and 14 were hypomethylated (no overlap with the males), out of 544,659 bins. Together, these results confirmed that maternal SMCHD1 as little to no impact on genome-wide DNA methylation at the morula stage.

### Absence of maternal SMCHD1 causes biallelic expression of *Xist* in the same cells, but silencing is restored by E3.5 in blastocysts

As our transcriptomic data had single-embryo (16-cell) but not single-cell resolution, we could not discriminate between the possibility that maternal and paternal *Xist* were co-expressed in the very same cells of female *Smchd1*^*mat*Δ^ embryos, or rather that each parental allele of *Xist* was monoallelically expressed in individual cells. To overcome this limitation, we performed allele-specific RNA-FISH in E2.75 embryos (Figure 3a). Labelling efficiency in allele-specific RNA-FISH is more variable than standard RNA FISH (***Harris et al., 2019***), and we estimated labelling efficiency to be above 50%, with some variability from embryo to embryo. The female wild-type embryos showed only paternal *Xist* expression in all labelled cells, as expected for imprinted XCI at the 16-cell stage (Figure 3b and d). By contrast in the *Smchd1*^*mat*Δ^ female embryos, *Xist* was expressed biallelically with maternal *Xist* detectable in a subset of cells (from 2 out of 16, up to 13 out 16 cells, Figure 3b and d). In males, we observed maternal *Xist* only in *Smchd1*^*mat*Δ^ embryos (Figure 2c and e), also in a subset of cells, which was consistent with the transcriptomic data. Therefore the detection of both paternal and maternal *Xist* in the female *Smchd1*^*mat*Δ^ transcriptomes was not due to an alternating pattern of mono-allelic expression, as seen in random X-chromosome inactivation after chromosome choice, but indeed due to the loss of imprinting of *Xist*. Penetrance was again only partial, as we did not observe biallelic expression in every cell of every embryo, and the proportion of cells with biallelic expression was variable between embryos.

**Figure 3.**
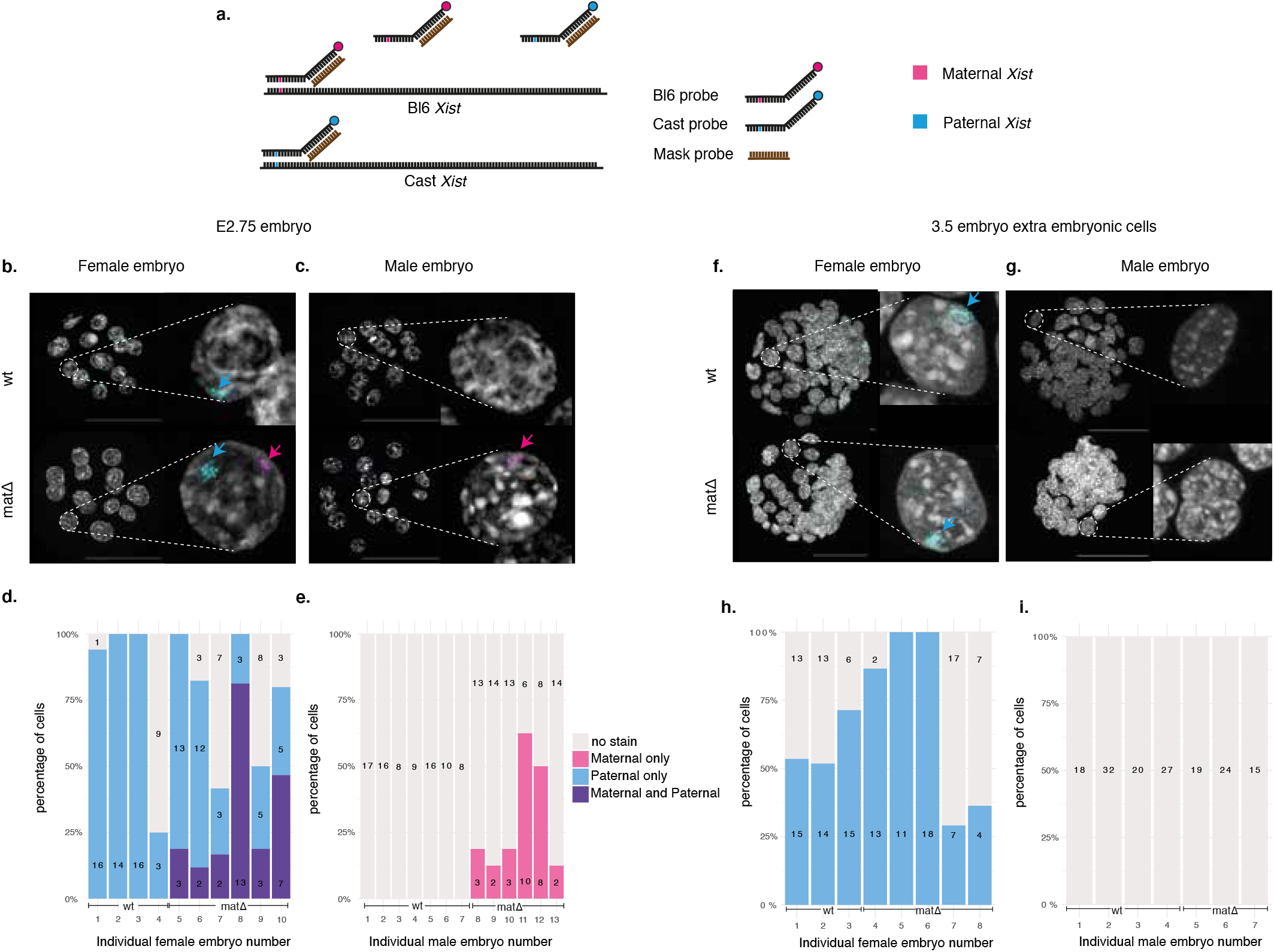
Maternal deletion of *Smchd1* results in transient biallelic *Xist* exist expression in morula. (a) Schematic representation of the allele-specific *Xist* RNA FISH. (b,c) Imaging of female (b) and (c) male wt and *Smchd1*^*mat*Δ^ E2.75 embryos. Maternal (BL6) and paternal (Cast) alleles are indicated by coloured arrows. (d,e) Percentage and number of cells in E2.75 female (d) and male (e) embryos with maternal, paternal or biallelic *Xist* expression. (f,g) Imaging of female (f) and (g) male wt and *Smchd1*^*mat*Δ^ E3.5 embryos. (h,i) Percentage and number of cells in E3.5 female (h) and male (i) embryos with maternal, paternal or biallelic *Xist* expression. Scale bar: 50 μm. Numbers of embryos and cells scored are indicated on the figure.

In a previous study where the *Xist* imprint was completely removed via *Eed* maternal deletion (***Harris et al., 2019***), biallelic *Xist* expression in early female embryos resolved into random X inactivation. By E3.5, a majority of female nuclei showed random monoalellic *Xist* expression (either paternal or maternal). Meanwhile male *Eed*^*mat*Δ^ embryos retained maternal *Xist* expression. To test whether the same was happening in *Smchd1*^*mat*Δ^ embryos, we performed allele-specific RNA-FISH on E3.5 embryos (early blastocysts, 32–64 cells). In the outer trophectoderm layer that gives rise to the placenta, *Xist* expression in *Smchd1*^*mat*Δ^ embryos was restored to the wild-type pattern: only paternal *Xist* expression in female embryos and no *Xist* expression in male embryos (Figure 3f-i). This contrasted with the *Eed*^*mat*Δ^ results. Although maternal *Xist* re-silencing in *Smchd1*^*mat*Δ^ female embryos may be explained by biased allele-choice because of the higher expression of paternal *Xist* over maternal *Xist*, counting and choice cannot explain the re-silencing of maternal *Xist* in male embryos. This instead suggest that the underlying imprint was successfully set up in oocyte development and maintained through the first 4–5 cell divisions in the absence of maternal SMCHD1. As restoration of imprinted expression aligns with the onset on zygotic expression of SMCHD1, we propose that zygotic SMCHD1 may rescue the loss of imprinted *Xist* expression. Regardless of the exact mechanism of imprint restoration, this places SMCHD1 downstream of the Polycomb-dependent imprint, similar to what we proposed for other non-canonically imprinted genes (***Wanigasuriya et al., 2020***).

### Maternal deletion of *Smchd1* does not affect *Xist* expression in E14.5 placentae but compromises XCI

Previously we showed loss of maternal SMCHD1 resulted in defects in the imprinted expression of some autosomal imprinted genes in the mid-gestation (E14.5) placenta, despite the presence of zygotic SMCHD1 for 11 days (***Wanigasuriya et al., 2020***). Although the correct pattern of imprinted *Xist* expression was restored by E3.5, we investigated whether any residual effects of maternal SMCHD1 ablation on imprinted X inactivation could be observed in E14.5 placentae.

We performed allele-specific bulk RNA-seq on the embryonic portion of female E14.5 placentae for matΔ, wild-type and heterozygous (paternally transmitted mutation) embryos. Comparing with heterozygous samples allowed to account for potential haploinsufficiency for *Smchd1* after zygotic SMCHD1 activation. Allelic *Xist* expression in *Smchd1*^*mat*Δ^ samples was indistinguishable from that of wild-type samples, consistent with the restored imprinted *Xist* expression by E3.5 (Figure 4a). Expression from the active X chromosome was also largely normal in heterozygous and *Smchd1*^*mat*Δ^ samples (Figure 4b, left panels). From the paternal inactive X however, 36 out of 179 informative genes were upregulated (informative: expressed and containing a SNP; 5% FDR) in heterozygous samples (Figure 4b, top right panel), while 107 out of 213 informative genes (5% FDR) were upregulated in the *Smchd1*^*mat*Δ^ samples (Figure 4b, bottom right panel). There were 34 X-linked genes that were upregulated in both genotypes (Figure 4c). These common genes tended to have larger log_2_ fold changes in the maternal null samples (p < 0.001, paired t-test, Figure 4d). This showed that while *Smchd1* haploinsufficiency impacted imprinted X inactivation in the E14.5 placentae, the loss of maternal SMCHD1 had a more severe effect, both in terms of the number of genes that escape silencing and the extent to which they escape.

**Figure 4.**
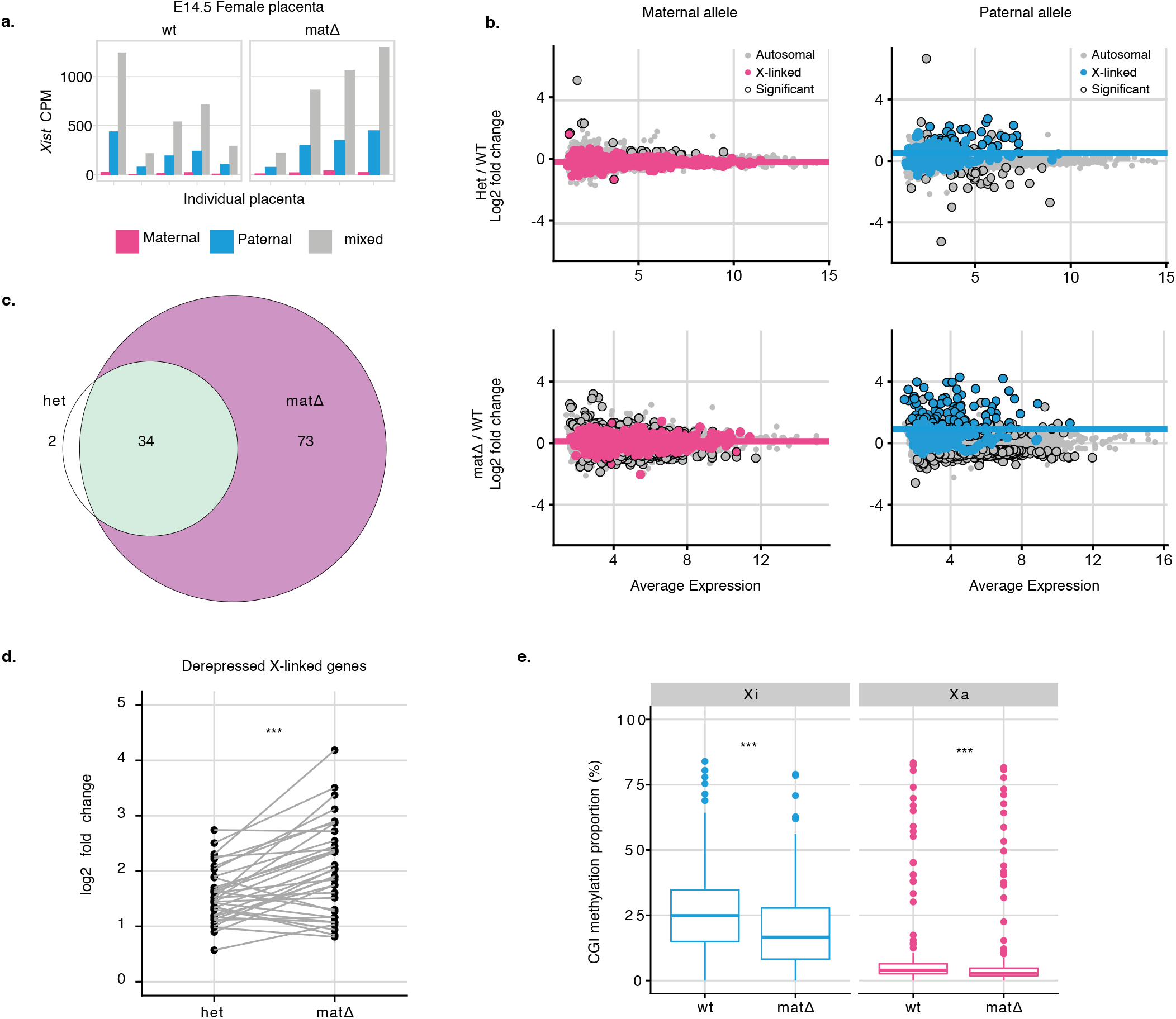
Maternal deletion of *Smchd1* results in failed silencing of the Xi in E14.5 placentae despite normal *Xist* expression. (a) *Xist* expression separated by maternal allele, paternal allele or total counts (without haplotyping, i.e. maternal + paternal + non-allele-specific reads) in female *Smchd1*^*mat*Δ^ or wt E14.5 placentae. The reads are shown as a proportion of the total library size (counts per million, CPM) before haplotyping. (b) Differential gene expression between *Smchd1*^*het*^ and *Smchd1*^*wt*^ E14.5 placentae, and *Smchd1*^*mat*Δ^ and Smchd1^*wt*^ in E14.5 placentae split by alleles. X-linked genes are coloured, differentially expressed genes that pass the genome-wide 5% FDR are circled. Average expression in log_2_ cpm. The paternal X is the inactive X is mouse placenta. Median log2-fold change of X-linked genes is plotted as a coloured horizontal line. (c) Overlap between X-linked genes that are significantly differentially expressed in *Smchd1*^*het*^ and in *Smchd1*^*mat*Δ^ placentae. (d) Comparison of the log_2_ fold changes of the differentially expressed paternal X-linked alleles common to the *Smchd1*^*het*^ and *Smchd1*^*mat*Δ^ placentae. p = 8e-5, paired t-test. (e) Distribution of CpG island methylation on the Xi and Xa in *Smchd1*^*mat*Δ^ and Smchd1^*wt*^ E14.5 female placentae. Xi: p < 1e-6; Xa: p = 2e-5; paired t-tests. n = 4 MMTV-Cre *Smchd1*^*mat*Δ^ and n = 5 wt; n = 6 het and littermate n = 4 wt control E14.5 placentae.

To investigate whether the failure to properly silence the inactive X could be linked to SMCHD1’s role in the methylation of CpG islands, we performed Reduced Representation Bisulfite Sequencing (RRBS) in *Smchd1*^*mat*Δ^ and wt female E14.5 placentae. CpG island methylation was reduced in *Smchd1*^*mat*Δ^ placentae, from 25% median methylation to 15% (Figure 4e). The low level of methylation observed in the placental tissue is as expected as this tissue has less methylation than embryonic tissues (***Schroeder et al., 2015***). The failure to silence Xi genes was thus correlated with a failure to adequately methylate the Xi CpG islands.

These data show that the perturbations induced by the lack of SMCHD1 preimplantation persist for at least 11 days post zygotic *Smchd1* activation, despite normal *Xist* expression at E3.5 and E14.5.

## Discussion

Previously we identified that SMCHD1 modulates the imprinted expression of a set of autosomal genes that we predicted was secondary to deposition of H3K27me3 by PRC2 (***Wanigasuriya et al., 2020***). This happened at two classes of loci: genes where the PRC2 mark is the primary imprint (non-canonical imprinted genes), and imprinted clusters where the primary DNA methylation imprint leads to secondary H3K27me3 deposition. Here we extend these findings, demonstrating that maternal SMCHD1 also enforces imprinted expression of the long non-coding RNA *Xist* during preimplantation embryo development. *Xist* belongs to the class of non-canonical imprinted genes, its promoter being labelled with H2K119ub and H3K27me3 during oocyte development (***Inoue et al., 2017; Mei et al., 2021; Chen et al., 2021***). In the absence of maternal SMCHD1, we observed biallelic *Xist* expression in female E2.75 embryos, and maternal *Xist* expression in male embryos.

Previous work by ***Harris et al. (2019***) and ***Inoue et al. (2017***) showed that loss of *Xist* imprinting leads to failed imprinted XCI. A maternal deletion of Polycomb gene *Eed* led to the erasure of the imprint on maternal *Xist* and complete loss of *Xist* imprinted expression. Harris et al. observed subsequent male lethality and a conversion from imprinted to random XCI in the female placentae. However, upon maternal *Smchd1* deletion we observed neither sex-specific embryonic lethality (***Wanigasuriya et al., 2020***) nor random XCI in the female placentae. By contrast, *Xist* loss of imprinting was incompletely penetrant (in a subset of embryos and cells) at E2.75 and normal maternal *Xist* silencing was fully restored by E3.5. We interpret the rescue of maternal *Xist* silencing, coinciding with zygotic SMCHD1 synthesis, as an indication that the underlying Polycomb imprint on *Xist* remained intact. This once again places SMCHD1 downstream of the Polycomb machinery. Recent work has shown that PRC1-deposited H2AK119ub is also involved in imprinted *Xist* expression and imprinted XCI (***Mei et al., 2021; Chen et al., 2021***). Since a PRC1-dependent model of SMCHD1 recruitment has been reported for the inactive X (***Jansz et al., 2018b; Wang et al., 2019***), this model likely extends to *Xist* and other non-canonical imprinted genes. Thus a single H2AK119ub-dependent recruitment mechanism could apply to both maternal and zygotic SMCHD1, at autosomes as well as at the X chromosome.

Although *Xist* loss of imprinting was only transient, the absence of SMCHD1 for the first three days of embryonic development had lasting effects on X inactivation. The initial phases of X inactivation, driven by the *Xist* long non-coding RNA, did not strictly require SMCHD1 to silence genes: *Xist* expression from the maternal allele was able to initiate gene silencing on the X chromosome in male E2.75 embryos lacking SMCHD1, and paternal X silencing was not abolished in E2.75 *Smchd1*^*mat*Δ^ female embryos. However, silencing efficiency was reduced in these females, which might be attributable to several factors. Biallelic *Xist* expression may delay the commitment to silencing or dilute silencing between the two X chromosomes. Alternatively, maternal SMCHD1 could contribute to some of the early paternal X silencing in addition to its role in imprinted maternal *Xist* repression. More surprisingly, X inactivation defects were observable in E14.5 female placentae, despite more than 10 days of zygotic SMCHD1 presence. Half of the detectable Xi-linked genes were not appropriately silenced, which could not simply be explained by *Smchd1* haploinsufficiency. In addition, failed gene silencing was associated with a failure to methylate CpG islands on the Xi to the same level as the wild-type. This persisting disruption to epigenetic silencing bore similarities with two other maternal effects seen in embryos with maternal *Smchd1* deletions: the partial loss of autosomal imprinting in the mid-gestation placentae (***Wanigasuriya et al., 2020***) and the disrupted *Hox* gene regulation in tissue of the embryo-proper (***Benetti et al., 2021***).

From this cumulative evidence, it is clear that maternal SMCHD1 is required during preimplantation development to set up an epigenetic state that is required for correct gene regulation later on. What this particular epigenetic state is and how SMCHD1 creates it remains obscure, but it is tempting to speculate that, like for SMCHD1 recruitment, a single mechanism explains SMCHD1’s mode of action for both maternal and zygotic SMCHD1, at all of its diverse targets. SMCHD1 is required for the repression of *Hox* genes, protocadherin clusters, imprinted genes, the inactive X, and tandem repeat arrays (***Jansz et al., 2018a; Benetti et al., 2021; Gendrel et al., 2012; Chen et al., 2015; Mould et al., 2013; Blewitt et al., 2008; Lemmers et al., 2012; Gdula et al., 2019***). All these targets display abundant Polycomb marks. In the absence of SMCHD1, H3K27me3 spreads and H3K9me3 is lost on the Xi (***Jansz et al., 2018b; Ichihara et al., 2021***). SMCHD1’s role at its targets may be to facilitate the positive feedback loops of Polycomb repression (***Blackledge and Klose, 2021***), concentrating the Polycomb machinery, Polycomb marks and repressors at specific loci to both reach a threshold required for efficient silencing as well as avoid ectopic redistribution of Polycomb. This mechanism would be conceptually close to what has been proposed for plant MORC proteins, GKHL ATPases like SMCHD1, which are proposed to anchor a chromatin silencing pathway to target loci (***Xue et al., 2021***). Ensuring focal enrichment of Polycomb repressive marks might in turn allow adjacent regions to adopt other chromatin states, explaining SMCHD1’s proposed role as an insulator (***Chen et al., 2015; Jansz et al., 2018a; Gdula et al., 2019***). These well-defined linear chromatin blocks would influence the three-dimensional self-organisation of the chromatin into domains of cognate epigenetic states, perhaps explaining the effect of SMCHD1 on long-range chromatin interactions (***Jansz et al., 2018a; Wang et al., 2018***). The potential role of SMCHD1 in solidifying initial silencing by Polycomb may then allow some of its targets to transition to other modes of repression, in particular H3K9 methylation and DNA methylation. Preimplantation Polycomb imprints acquire secondary DNA methylation and H3K9me2 in the placenta (***Hanna et al., 2019; Chen et al., 2019; Zeng et al., 2021; Andergassen et al., 2021; Raas et al., 2021***), similar to the Xi CpG islands becoming methylated by DNMT3B and H3K9me3 accumulating on the Xi (***Gendrel et al., 2012; Keniry et al., 2016; Ichihara et al., 2021***). In the absence of SMCHD1, these transitions fail: DNA methylation at non-canonical imprinted gene *Jade1* does not accumulate (***Wanigasuriya et al., 2020***), nor does Xi CpG island methylation (***Gendrel et al., 2012***)(this study).

SMCHD1 links together epigenetic processes that appear more and more closely related as our knowledge increases. Non-canonical imprinting, imprinted X inactivation, aspects of canonical imprinting and random X inactivation all borrow from the same Polycomb/SMCHD1/H3K9me/DNA methylation toolbox. Each process offers a window into a general but complex interplay of epigenetic mechanisms. Further elucidation of SMCHD1’s molecular mechanisms will shed light on all these fundamental processes.

## Materials and Methods

### Mouse genetics

All mice were bred and maintained in house at The Walter and Eliza Hall Institute of Medical Research (WEHI) using procedures approved by in-house ethics approval committee (approval numbers 2014.026, 2018.004, 2020.048 and 2020.050).

*Smchd1*^*mat*Δ^ embryos were produced from a cross between *Smchd1*^*-/fl*^ *MMTV-Cre*^*T/+*^ dams and CAST/EiJ sires. *Smchd1*^*het*^ embryos were produced from the reciprocal cross. Control *Smchd1*^*wt*^ embryos were produced from crosses between *Smchd1*^*fl/fl*^ *MMTV-Cre*^*+/+*^ dams and CAST/EiJ sires, and as littermates of the *Smchd1*^*het*^ embryos.

The MMTV-Cre *Smchd1*^*-/fl*^ line was generated by backcrossing MMTV-Cre transgene line A (***Wagner et al., 1997***) onto the C57BL/6 background from the FVB/N background for more than 10 generations. These mice contain a combination of *Smchd1* deleted allele (*Smchd1*^*-*^) in trans to the *Smchd1* floxed (*Smchd1*^fl^) allele (***de Greef et al., 2018***). The CAST/EiJ strain used to achieve polymorphism necessary for allele-specific analyses was purchased from the Jackson laboratories.

### Single-embryo methylome and transcriptome sequencing

Embryo collection, library preparation and data preprocessing were as described in ***Wanigasuriya et al. (2020***). Male and female embryos were analysed in the same way.

RNA-seq reads from the E2.75 embryos were trimmed for adapter and low quality sequences using TrimGalore! v0.4.4, before mapping onto the GRCm38 mouse genome reference N-masked for Cast SNPs prepared with SNPsplit v0.3.2 (***Krueger and Andrews, 2016***) with HISAT2 v2.0.5 (***Kim et al., 2015***), in paired-end mode and disabling soft-clipping. Gene counts were obtained in R 3.5.1 (***R Core Team, 2019***) from bam files with the featureCounts function from the Rsubread package v1.32.1 (***Liao et al., 2014, 2019***), provided with the GRCm38.90 GTF annotation downloaded from Ensembl, and ignoring multi-mapping or multi-overlapping reads. Lowly expressed genes were filtered out with the filterByExpr function with option min.prop = 0.33 in edgeR v3.24.0 (***Robinson et al., 2010; McCarthy et al., 2012***). Gene counts were normalised with the TMM method (***Robinson and Oshlack, 2010***). Differential gene expression between the *Smchd1*^*mat*Δ^ and Smchd1^*wt*^ embryos was performed using the glmFit and glmLRT functions. P-values were corrected with the Benjamini-Hochberg method (***Benjamini and Hochberg, 1995***). Differential expression results were visualized with Glimma 2.2.0 (***Su et al., 2017; Kariyawasam et al., 2021***).

Whole-genome bisulfite analysis of single E2.75 embryos was performed as in (***Wanigasuriya et al., 2020***).

### Bulk RNA-seq

RNA-seq libraries from the embryonic portion of E14.5 mouse placentae were made as described in ***Wanigasuriya et al. (2020***). Differential expression analysis was performed using the same strategy as for the above single-embryo RNA-seq.

### RRBS

Library preparation and analysis was identical to ***Wanigasuriya et al. (2020***).

### Allele-specific RNA-FISH on preimplantation embryos

#### Probe preparation

Allele-specific *Xist* RNA FISH probes were generated as described (***Levesque et al., 2013***). Briefly, a set of short oligonucleotide probes (5 probes for each *Xist* allele) were designed to uniquely detect either the Bl6 or the Cast alleles of *Xist* exon 7. Each probe contained single nucleotide polymorphism (SNP) located at the fifth base pair position from the 5’ end that differs between the Bl6 and Cast. The 3’ end of each oligonucleotide probe was fluorescently tagged using Quasar dyes (Biosearch technologies). Bl6-specific oligos were labelled with Quasar 570 and Cast-specific oligos labelled with Quasar 670. In addition to labeled SNP-overlapping oligonucleotides, a panel of 5 ‘mask’ oligonucleotides were also synthesized (IDT). Exon 7 of *Xist* RNA was selected as the strand-specific*Xist* guide probe. Exon 7 specific primers were designed (IDT) with T3 and T7 promoter overhangs. Exon 7 was amplified from 50 ng of an *Xist* cDNA clone (***Wutz and Jaenisch, 2000***). Briefly, the PCR reaction contained cDNA, 5x Phusion HF reaction buffer (Cat # 13058S), 1 μL Phusion Taq, 10 mM dNTP, and 10 μM per forward and reverse primers. PCR cycle conditions were 98°C for 2 minutesutes; 30 cycles of 98°C for 30 seconds, 58°C for 30 seconds and 72°C for 30 seconds; 72°C for 4 minutes. PCR product was isolated using QIAquick gel extraction kit (Qiagen) according to manufacturer’s instructions. Strand-specific *Xist* RNA probe was labelled with Fluorescein-12-UTP (Roche, Cat # 11427857910) and ethanol precipitated as previously described (***Hinten et al., 2016***). Probe was re-suspended in hybridisation buffer containing 10% dextran sulfate, 2X saline-sodium citrate (SSC) and 10% formamide.

#### Allele-specific RNA FISH

E2.75 embryos were collected and the zona pellucida removed by keeping in acid tyrode’s solution (Sigma) for 2 minutes. Embryos were placed in the middle of Denhardt’s treated cover slips in 1x PBS 6 mg/ml BSA using finely pulled Pasteur pipette. Excess 1x PBS 6 mg/ml BSA was aspirated and embryos let dry for 20–30 minutes. Embryos were fixed and permeabilised with 50 μL of 1% PFA in 1x PBS with 0.05% Tergitol for 5 minutes. Embryos were rinsed with three changes of 70% ethanol then dehydrated through an ethanol series (85%, 95%,100%) 2 minutes each at room temperature. Samples were then air-dried for 5–10 minutes.

Allele-specific *Xist* RNA FISH was performed on these embryos as previously described (***Harris et al., 2019***). The precipitated guide RNA probe was mixed with the Bl6 and Cast detection probes, to a final concentration of 5 nM per allele-specific oligo, and 10 nMmask probe, yielding a 1:1 mask:detection oligonucleotide ratio. Cover slips were hybridised to the combined probe overnight in a humid chamber at 37°C. After overnight hybridisation, samples were washed twice in 2x SSC with 10% formamide at 37°C for 30 minutes, followed by one wash in 2X SSC for five minutes at room temperature. A 1/250,000 dilution of DAPI (Invitrogen, Cat # D21490) was added to the second 2X SSC with 10% formamide wash. Cover slips were then mounted on slides in Vectashield (Vector Labs, Cat # H-1000). Stained samples were imaged immediately using a LSM 880 (Zeiss) microscope.

## Acknowledgments

We thank Andrea Morcom for assisting with preimplantation embryo flushing and Jessica Martin for mouse husbandry assistance. We also thank Sundeep Kalantry and Clair Harris for their assistance with allele specific FISH protocol.

## Authors’ contributions

IW: Conceptualization, Investigation, Data curation, Formal analysis, Methodology, Writing - original draft, Writing - review and editing; SAK, TB, EAR, KB: Investigation; HJL, AK: Methodology, Resources, Supervision; MER: Methodology, Resources, Supervision, Writing - review and editing; MEB: Con-ceptualization, Methodology, Resources, Supervision, Writing - original draft, Writing - review and editing; QG: Conceptualization, Investigation, Data curation, Formal analysis, Methodology, Supervision, Writing - original draft, Writing - review and editing;

## Data availability

Female single-embryo Methylome and Transcriptome sequencing, female E14.5 placenta RRBS and RNA-seq raw and processed data are available under GEO accession GSE186315. Male data from ***Wanigasuriya et al***. (***2020***) are available under BioProject accession PRJNA530651.

## Appendix 1

### Supplementary files

1. tables of differential expression results: male and female E2.75 mat Δvs wt embryos (total and allelic), female E14.5 mat Δvs wt placentae (allelic), female E14.5 het vs wt placentae (allelic).
2. tables of differential methylation results: female E2.75 mat Δvs wt embryos in 10 kb windows, at promoters and at CGIs.

**Figure 1–Figure supplement 1.**
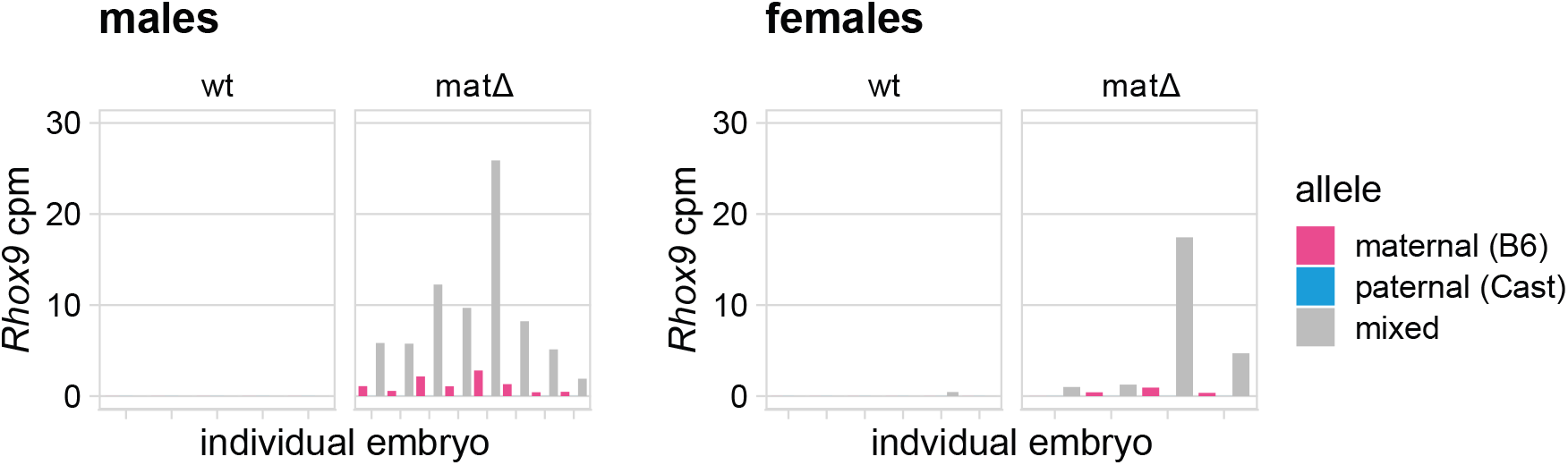
*Rhox9* expression in individual male and female wt and *Smchd1*^*mat*Δ^ E2.75 embryos. cpm: counts per million (of total library size before haplotyping). “Mixed” counts refer to counts without haplotyping. Females: n = 6 wt and 4 matΔ; males: n = 5 wt and 8 matΔ.

**Figure 1–Figure supplement 2.**
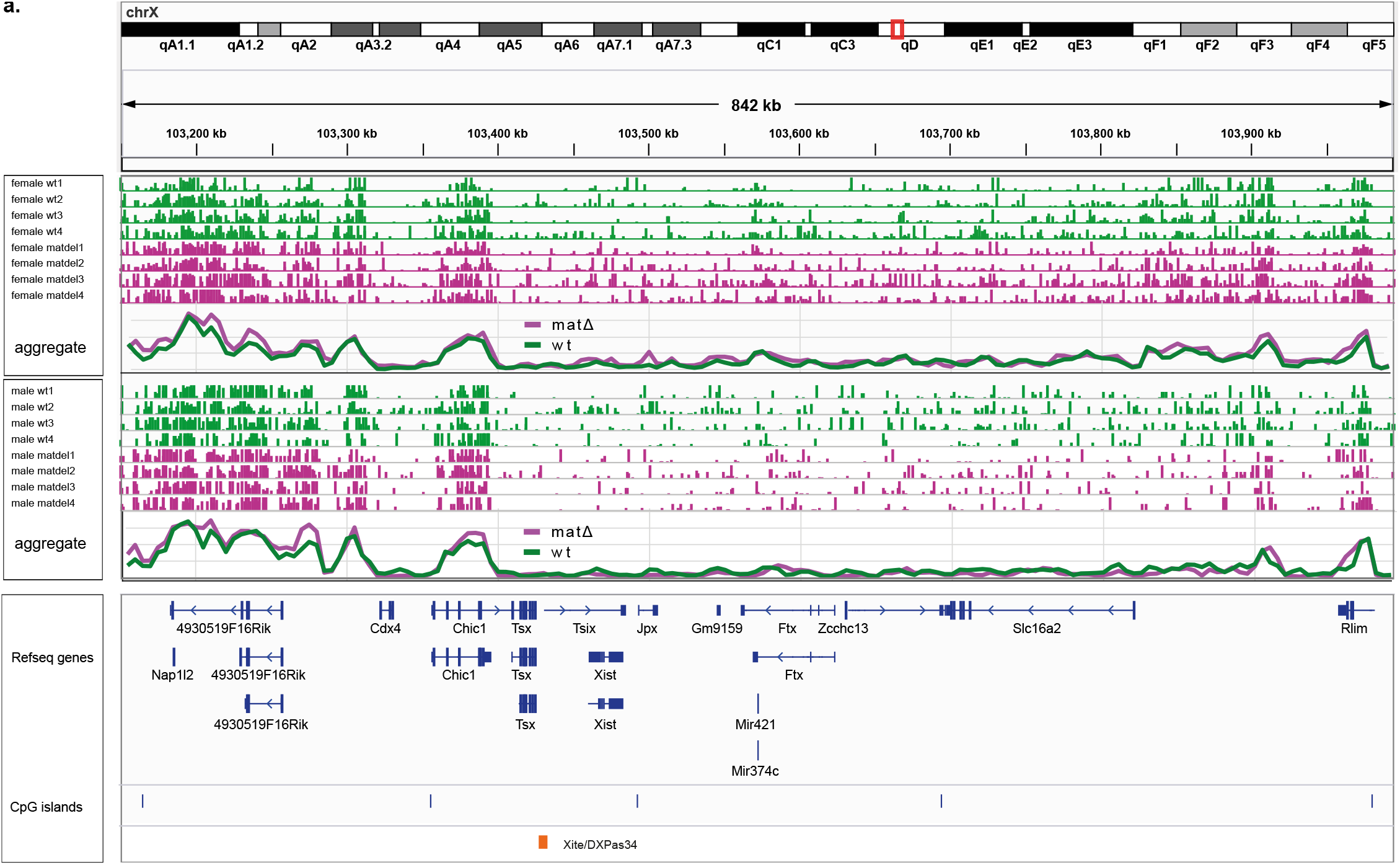
DNA methylation at the X inactivation center in female and male E2.75 wt embryos and *Smchd1*^*mat*Δ^ embryos. Histogram tracks show methylation in single embryos at individual CpG sites (0–100%, 4 wild-type and 4 mat embryos shown for each sex). Note that coverage in single-embryo whole-genome bisulfite sequencing is sparse, only 0–2X. The aggregate line plots show the average methylation per genotype across 10 kb windows (sliding by 5 kb, 0–100%). Females: n = 6 wt and 4 matΔ; males: n = 5 wt and 8 matΔ.

**Figure 2–Figure supplement 1.**
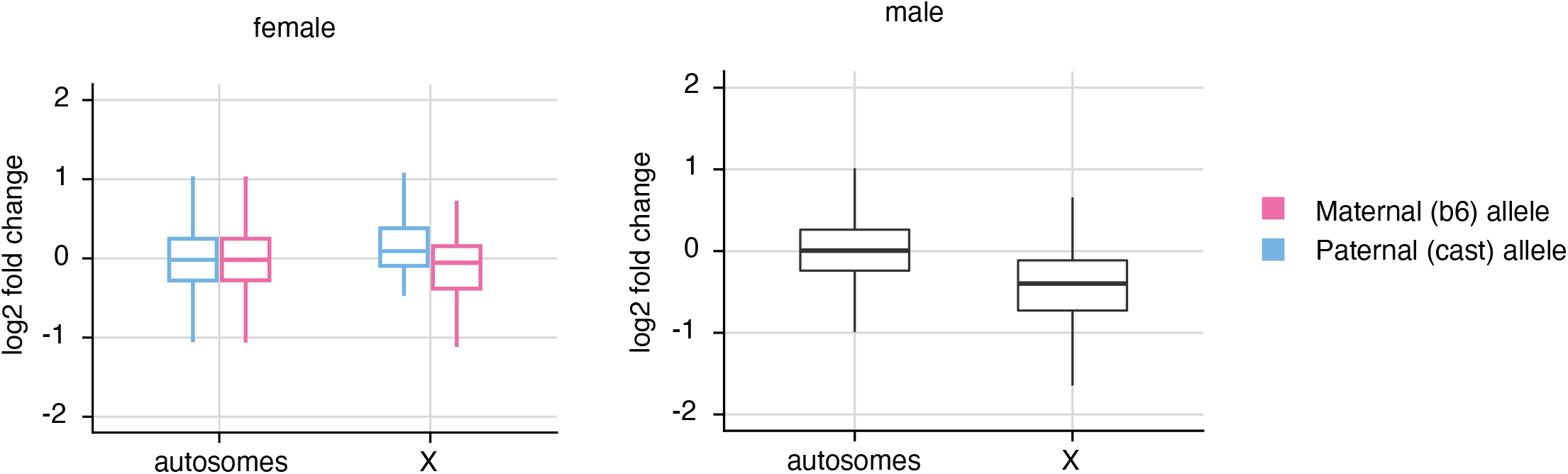
Distribution of *Smchd1*^*mat*Δ^ vs wt gene expression log2 fold changes on autosomes and the X chromosome for male and female E2.75 embryos, retaining only matΔembryos with loss of *Xist* imprinting.

**Figure 2–Figure supplement 2.**
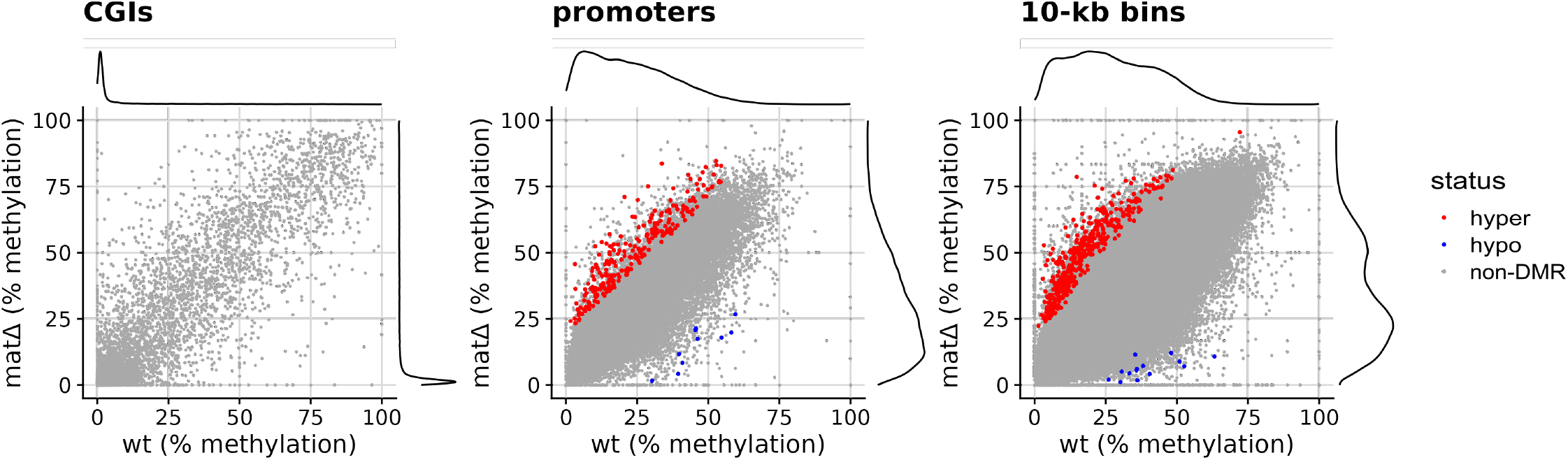
Whole-genome differential methylation analysis between female *Smchd1*^*mat*Δ^ and wild-type E2.75 embryos. For CpG islands (CGIs, 13k regions), promoters (−4 kb to +1 kb regions, 52k regions) and 10-kb windows (sliding by 5 kb, 500k regions), the average methylation level in wild types is plotted against the average methylation in *Smchd1*^*mat*Δ^ embryos. Significant Differentially Methylated Regions (DMRs, FDR < 5% and absolute difference in methylation > 20%) are coloured in red (hypermethylation) or blue (hypomethylation). Females: n = 6 wt and 4 matΔ.

## References

Andergassen D, Smith ZD, Rinn JL, Meissner A. Diverse mechanisms for epigenetic imprinting in mammals. bioRxiv. 2021 Apr; http://biorxiv.org/lookup/doi/10.1101/2021.04.30.442087, doi: 10.1101/2021.04.30.442087.

Benetti N, Gouil Q, del Fierro AT, Beck T, Breslin K, Keniry A, McGlinn E, Blewitt ME. Maternal SMCHD1 regulates Hox gene expression and patterning in the mouse embryo. bioRxiv. 2021 Sep; http://biorxiv.org/lookup/doi/10.1101/2021.09.08.459528, doi: 10.1101/2021.09.08.459528.

Benjamini Y, Hochberg Y. Controlling the False Discovery Rate: A Practical and Powerful Approach to Multiple Testing. Journal of the Royal Statistical Society: Series B (Methodological). 1995 Jan; 57(1):289–300. https://onlinelibrary.wiley.com/doi/10.1111/j.2517-6161.1995.tb02031.x, doi: 10.1111/j.2517-6161.1995.tb02031.x.

Berletch JB, Deng X, Nguyen DK, Disteche CM. Female Bias in Rhox6 and 9 Regulation by the Histone Demethylase KDM6A. PLoS Genetics. 2013 May; 9(5):e1003489. https://dx.plos.org/10.1371/journal.pgen.1003489, doi: 10.1371/journal.pgen.1003489.

Blackledge NP, Klose RJ. The molecular principles of gene regulation by Polycomb repressive complexes. Nature Reviews Molecular Cell Biology. 2021 Aug; https://www.nature.com/articles/s41580-021-00398-y, doi: 10.1038/s41580-021-00398-y.

Blewitt ME, Gendrel AV, Pang Z, Sparrow DB, Whitelaw N, Craig JM, Apedaile A, Hilton DJ, Dunwoodie SL, Brockdorff N, Kay GF, Whitelaw E. SmcHD1, containing a structural-maintenance-of-chromosomes hinge domain, has a critical role in X inactivation. Nature Genetics. 2008 May; 40(5):663–669. http://www.nature.com/articles/ng.142, doi: 10.1038/ng.142.

Borensztein M, Syx L, Ancelin K, Diabangouaya P, Picard C, Liu T, Liang JB, Vassilev I, Galupa R, Servant N, Barillot E, Surani A, Chen CJ, Heard E. Xist-dependent imprinted X inactivation and the early developmental consequences of its failure. Nature Structural & Molecular Biology. 2017 Mar; 24(3):226–233. http://www.nature.com/articles/nsmb.3365, doi: 10.1038/nsmb.3365.

Chen K, Hu J, Moore DL, Liu R, Kessans SA, Breslin K, Lucet IS, Keniry A, Leong HS, Parish CL, Hilton DJ, RJLF Lemmers, van der Maarel SM, Czabotar PE, Dobson RCJ, Ritchie ME, Kay GF, Murphy JM, Blewitt ME. Genome-wide binding and mechanistic analyses of Smchd1-mediated epigenetic regulation. Proceedings of the National Academy of Sciences. 2015 Jul; 112(27):E3535–E3544. http://www.pnas.org/lookup/doi/10.1073/pnas.1504232112, doi: 10.1073/pnas.1504232112.

Chen Z, Djekidel MN, Zhang Y. Distinct dynamics and functions of H2AK119ub1 and H3K27me3 in mouse preimplantation embryos. Nature Genetics. 2021 Apr; http://www.nature.com/articles/s41588-021-00821-2, doi: 10.1038/s41588-021-00821-2.

Chen Z, Yin Q, Inoue A, Zhang C, Zhang Y. Allelic H3K27me3 to allelic DNA methylation switch maintains noncanonical imprinting in extraembryonic cells. Science Advances. 2019 Dec; 5(12):eaay7246. https://advances.sciencemag.org/lookup/doi/10.1126/sciadv.aay7246, doi: 10.1126/sciadv.aay7246.

Chiba H, Hirasawa R, Kaneda M, Amakawa Y, Li E, Sado T, Sasaki H. De novo DNA methylation independent establishment of maternal imprint on X chromosome in mouse oocytes. genesis. 2008 Dec; 46(12):768–774. https://onlinelibrary.wiley.com/doi/10.1002/dvg.20438, doi: 10.1002/dvg.20438.

Cooper DW, Vandeberg JL, Sharman GB, Poole WE. Phosphoglycerate Kinase Polymorphism in Kangaroos provides Further Evidence for Paternal X Inactivation. Nature New Biology. 1971 Mar; 230(13):155–157. http://www.nature.com/articles/newbio230155a0, doi: 10.1038/newbio230155a0.

Deakin JE, Chaumeil J, Hore TA, Marshall Graves JA. Unravelling the evolutionary origins of X chromosome inactivation in mammals: insights from marsupials and monotremes. Chromosome Research. 2009 Jul; 17(5):671–685. http://link.springer.com/10.1007/s10577-009-9058-6, doi: 10.1007/s10577-009-9058-6.

Galupa R, Heard E. X-Chromosome Inactivation: A Crossroads Between Chromosome Architecture and Gene Regulation. Annual Review of Genetics. 2018 Nov; 52(1):535–566. https://www.annualreviews.org/doi/10.1146/annurev-genet-120116-024611, doi: 10.1146/annurev-genet-120116-024611.

Gdula MR, Nesterova TB, Pintacuda G, Godwin J, Zhan Y, Ozadam H, McClellan M, Moralli D, Krueger F, Green CM, Reik W, Kriaucionis S, Heard E, Dekker J, Brockdorff N. The non-canonical SMC protein SmcHD1 antagonises TAD formation and compartmentalisation on the inactive X chromosome. Nature Communications. 2019 Dec; 10(1):30. http://www.nature.com/articles/s41467-018-07907-2, doi: 10.1038/s41467-018-07907-2.

Gendrel AV, Apedaile A, Coker H, Termanis A, Zvetkova I, Godwin J, Tang YA, Huntley D, Montana G, Taylor S, Giannoulatou E, Heard E, Stancheva I, Brockdorff N. Smchd1-Dependent and -Independent Pathways Determine Developmental Dynamics of CpG Island Methylation on the Inactive X Chromosome. Developmental Cell. 2012 Aug; 23(2):265–279. https://linkinghub.elsevier.com/retrieve/pii/S1534580712002857, doi: 10.1016/j.devcel.2012.06.011.

de Greef JC, Krom YD, den Hamer B, Snider L, Hiramuki Y, van den Akker RFP, Breslin K, Pakusch M, Salvatori DCF, Slütter B, Tawil R, Blewitt ME, Tapscott SJ, van der Maarel SM. Smchd1 haploinsufficiency exacerbates the phenotype of a transgenic FSHD1 mouse model. Human Molecular Genetics. 2018 Feb; 27(4):716–731. https://academic.oup.com/hmg/article/27/4/716/4769583, doi: 10.1093/hmg/ddx437.

Hanna CW, Pérez-Palacios R, Gahurova L, Schubert M, Krueger F, Biggins L, Andrews S, Colomé-Tatché M, Bourc’his D, Dean W, Kelsey G. Endogenous retroviral insertions drive non-canonical imprinting in extra-embryonic tissues. Genome Biology. 2019 Dec; 20(1):225. https://genomebiology.biomedcentral.com/articles/10.1186/s13059-019-1833-x, doi: 10.1186/s13059-019-1833-x.

Harris C, Cloutier M, Trotter M, Hinten M, Gayen S, Du Z, Xie W, Kalantry S. Conversion of random X-inactivation to imprinted X-inactivation by maternal PRC2. eLife. 2019 Apr; 8:e44258. https://elifesciences.org/articles/44258, doi: 10.7554/eLife.44258.

Heard E, Chaumeil J, Masui O, Okamoto I. Mammalian X-Chromosome Inactivation: An Epigenetics Paradigm. Cold Spring Harbor Symposia on Quantitative Biology. 2004 Jan; 69(0):89–102. http://symposium.cshlp.org/cgi/doi/10.1101/sqb.2004.69.89, doi: 10.1101/sqb.2004.69.89.

Hinten M, Maclary E, Gayen S, Harris C, Kalantry S. Visualizing Long Noncoding RNAs on Chromatin. In: Feng Y, Zhang L, editors. Long Non-Coding RNAs, vol. 1402 New York, NY: Springer New York; 2016.p. 147–164. http://link.springer.com/10.1007/978-1-4939-3378-5_12, doi: 10.1007/978-1-4939-3378-5_12, series Title: Methods in Molecular Biology.

Ichihara S, Nagao K, Sakaguchi T, Obuse C, Sado T. SmcHD1 underlies the formation of H3K9me3 blocks on the inactive X chromosome in mice. bioRxiv. 2021 Aug; http://biorxiv.org/lookup/doi/10.1101/2021.08.23.457321, doi: 10.1101/2021.08.23.457321.

Inoue A, Jiang L, Lu F, Zhang Y. Genomic imprinting of Xist by maternal H3K27me3. Genes & Development. 2017 Oct; 31(19):1927–1932. http://genesdev.cshlp.org/lookup/doi/10.1101/gad.304113.117, doi: 10.1101/gad.304113.117.

Jansz N, Keniry A, Trussart M, Bildsoe H, Beck T, Tonks ID, Mould AW, Hickey P, Breslin K, Iminitoff M, Ritchie ME, McGlinn E, Kay GF, Murphy JM, Blewitt ME. Smchd1 regulates long-range chromatin interactions on the inactive X chromosome and at Hox clusters. Nature Structural & Molecular Biology. 2018 Sep; 25(9):766–777. http://www.nature.com/articles/s41594-018-0111-z, doi: 10.1038/s41594-018-0111-z.

Jansz N, Nesterova T, Keniry A, Iminitoff M, Hickey PF, Pintacuda G, Masui O, Kobelke S, Geoghegan N, Breslin KA, Willson TA, Rogers K, Kay GF, Fox AH, Koseki H, Brockdorff N, Murphy JM, Blewitt ME. Smchd1 Targeting to the Inactive X Is Dependent on the Xist-HnrnpK-PRC1 Pathway. Cell Reports. 2018 Nov; 25(7):1912–1923.e9. https://linkinghub.elsevier.com/retrieve/pii/S2211124718316334, doi: 10.1016/j.celrep.2018.10.044.

Kariyawasam H, Su S, Voogd O, Ritchie ME, Law CW. Dashboard-style interactive plots for RNA-seq analysis are R Markdown ready with Glimma 2.0. bioRxiv. 2021 Aug; http://biorxiv.org/lookup/doi/10.1101/2021.07.30.454464, doi: 10.1101/2021.07.30.454464.

Keniry A, Gearing LJ, Jansz N, Liu J, Holik AZ, Hickey PF, Kinkel SA, Moore DL, Breslin K, Chen K, Liu R, Phillips C, Pakusch M, Biben C, Sheridan JM, Kile BT, Carmichael C, Ritchie ME, Hilton DJ, Blewitt ME. Setdb1-mediated H3K9 methylation is enriched on the inactive X and plays a role in its epigenetic silencing. Epigenetics & Chromatin. 2016 Dec; 9(1):16. https://epigeneticsandchromatin.biomedcentral.com/articles/10.1186/s13072-016-0064-6, doi: 10.1186/s13072-016-0064-6.

Kim D, Langmead B, Salzberg SL. HISAT: a fast spliced aligner with low memory requirements. Nature Methods. 2015 Apr; 12(4):357–360. http://www.nature.com/articles/nmeth.3317, doi: 10.1038/nmeth.3317.

Krueger F, Andrews SR. SNPsplit: Allele-specific splitting of alignments between genomes with known SNP genotypes. F1000Research. 2016 Jun; 5:1479. https://f1000research.com/articles/5-1479/v1, doi: 10.12688/f1000research.9037.1.

RJLF Lemmers, Tawil R, Petek LM, Balog J, Block GJ, Santen GWE, Amell AM, van der Vliet PJ, Almomani R, Straasheijm KR, Krom YD, Klooster R, Sun Y, den Dunnen JT, Helmer Q, Donlin-Smith CM, Padberg GW, van Engelen BGM, de Greef JC, Aartsma-Rus AM, et al. Digenic inheritance of an SMCHD1 mutation and an FSHD-permissive D4Z4 allele causes facioscapulohumeral muscular dystrophy type 2. Nature Genetics. 2012 Dec; 44(12):1370–1374. http://www.nature.com/articles/ng.2454, doi: 10.1038/ng.2454.

Levesque MJ, Ginart P, Wei Y, Raj A. Visualizing SNVs to quantify allele-specific expression in single cells. Nature Methods. 2013 Sep; 10(9):865–867. http://www.nature.com/articles/nmeth.2589, doi: 10.1038/nmeth.2589.

Liao Y, Smyth GK, Shi W. featureCounts: an efficient general purpose program for assigning sequence reads to genomic features. Bioinformatics. 2014 Apr; 30(7):923–930. https://academic.oup.com/bioinformatics/article-lookup/doi/10.1093/bioinformatics/btt656, doi: 10.1093/bioinformatics/btt656.

Liao Y, Smyth GK, Shi W. The R package Rsubread is easier, faster, cheaper and better for alignment and quantification of RNA sequencing reads. Nucleic Acids Research. 2019 May; 47(8):e47–e47. https://academic.oup.com/nar/article/47/8/e47/5345150, doi: 10.1093/nar/gkz114.

Lyon MF. Sex chromatin and gene action in the mammalian X-chromosome. American Journal of Human Genetics. 1962 Jun; 14:135–148.

Lyon MF. Gene Action in the X-chromosome of the Mouse (Mus musculus L.). Nature. 1961 Apr; 190(4773):372–373. http://www.nature.com/articles/190372a0, doi: 10.1038/190372a0.

Lyon MF. Attempts to test the inactive-X theory of dosage compensation in mammals. Genetical Research. 1963 Feb; 4(1):93–103. https://www.cambridge.org/core/product/identifier/S0016672300003451/type/journal_article, doi: 10.1017/S0016672300003451.

MacLean JA, Bettegowda A, Kim BJ, Lou CH, Yang SM, Bhardwaj A, Shanker S, Hu Z, Fan Y, Eckardt S, McLaughlin KJ, Skoultchi AI, Wilkinson MF. The Rhox Homeobox Gene Cluster Is Imprinted and Selectively Targeted for Regulation by Histone H1 and DNA Methylation. Molecular and Cellular Biology. 2011 Mar; 31(6):1275–1287. https://journals.asm.org/doi/10.1128/MCB.00734-10, doi: 10.1128/MCB.00734-10.

McCarthy DJ, Chen Y, Smyth GK. Differential expression analysis of multifactor RNA-Seq experiments with respect to biological variation. Nucleic Acids Research. 2012 May; 40(10):4288–4297. https://academic.oup.com/nar/article/40/10/4288/2411520, doi: 10.1093/nar/gks042.

McGraw S, Oakes CC, Martel J, Cirio MC, de Zeeuw P, Mak W, Plass C, Bartolomei MS, Chaillet JR, Trasler JM. Loss of DNMT1o Disrupts Imprinted X Chromosome Inactivation and Accentuates Placental Defects in Females. PLoS Genetics. 2013 Nov; 9(11):e1003873. https://dx.plos.org/10.1371/journal.pgen.1003873, doi: 10.1371/journal.pgen.1003873.

Mei H, Kozuka C, Hayashi R, Kumon M, Koseki H, Inoue A. H2AK119ub1 guides maternal inheritance and zygotic deposition of H3K27me3 in mouse embryos. Nature Genetics. 2021 Apr; http://www.nature.com/articles/s41588-021-00820-3, doi: 10.1038/s41588-021-00820-3.

Mould AW, Pang Z, Pakusch M, Tonks ID, Stark M, Carrie D, Mukhopadhyay P, Seidel A, Ellis JJ, Deakin J, Wakefield MJ, Krause L, Blewitt ME, Kay GF. Smchd1 regulates a subset of autosomal genes subject to monoallelic expression in addition to being critical for X inactivation. Epigenetics & Chromatin. 2013 Dec; 6(1):19. https://epigeneticsandchromatin.biomedcentral.com/articles/10.1186/1756-8935-6-19, doi: 10.1186/1756-8935-6-19.

Okamoto I, Heard E. The dynamics of imprinted X inactivation during preimplantation development in mice. Cytogenetic and Genome Research. 2006; 113(1-4):318–324. https://www.karger.com/Article/FullText/90848, doi: 10.1159/000090848.

Okamoto I, Patrat C, Thépot D, Peynot N, Fauque P, Daniel N, Diabangouaya P, Wolf JP, Renard JP, Duranthon V, Heard E. Eutherian mammals use diverse strategies to initiate X-chromosome inactivation during development. Nature. 2011 Apr; 472(7343):370–374. http://www.nature.com/articles/nature09872, doi: 10.1038/nature09872.

Patrat C, Okamoto I, Diabangouaya P, Vialon V, Le Baccon P, Chow J, Heard E. Dynamic changes in paternal X-chromosome activity during imprinted X-chromosome inactivation in mice. Proceedings of the National Academy of Sciences. 2009 Mar; 106(13):5198–5203. http://www.pnas.org/cgi/doi/10.1073/pnas.0810683106, doi: 10.1073/pnas.0810683106.

Prissette M. Methylation profiles of DXPas34 during the onset of X-inactivation. Human Molecular Genetics. 2001 Jan; 10(1):31–38. https://academic.oup.com/hmg/article-lookup/doi/10.1093/hmg/10.1.31, doi: 10.1093/hmg/10.1.31.

R Core Team. R: A Language and Environment for Statistical Computing. R Foundation for Statistical Computing, Vienna, Austria; 2019, https://www.R-project.org/.

Raas MWD, Zijlmans DW, Vermeulen M, Marks H. There is another: H3K27me3-mediated genomic imprinting. Trends in Genetics. 2021 Jul; p. S0168952521001876. https://linkinghub.elsevier.com/retrieve/pii/S0168952521001876, doi: 10.1016/j.tig.2021.06.017.

Reik W, Lewis A. Co-evolution of X-chromosome inactivation and imprinting in mammals. Nature Reviews Genetics. 2005 May; 6(5):403–410. http://www.nature.com/articles/nrg1602, doi: 10.1038/nrg1602.

Robinson MD, McCarthy DJ, Smyth GK. edgeR: a Bioconductor package for differential expression analysis of digital gene expression data. Bioinformatics. 2010 Jan; 26(1):139–140. https://academic.oup.com/bioinformatics/article-lookup/doi/10.1093/bioinformatics/btp616, doi: 10.1093/bioinformatics/btp616.

Robinson MD, Oshlack A. A scaling normalization method for differential expression analysis of RNA-seq data. Genome Biology. 2010; 11(3):R25. http://genomebiology.biomedcentral.com/articles/10.1186/gb-2010-11-3-r25, doi: 10.1186/gb-2010-11-3-r25.

Schroeder DI, Jayashankar K, Douglas KC, Thirkill TL, York D, Dickinson PJ, Williams LE, Samollow PB, Ross PJ, Bannasch DL, Douglas GC, LaSalle JM. Early Developmental and Evolutionary Origins of Gene Body DNA Methylation Patterns in Mammalian Placentas. PLOS Genetics. 2015 Aug; 11(8):e1005442. https://dx.plos.org/10.1371/journal.pgen.1005442, doi: 10.1371/journal.pgen.1005442.

Su S, Law CW, Ah-Cann C, Asselin-Labat ML, Blewitt ME, Ritchie ME. Glimma: interactive graphics for gene expression analysis. Bioinformatics. 2017 Jul; 33(13):2050–2052. https://academic.oup.com/bioinformatics/article/33/13/2050/2997287, doi: 10.1093/bioinformatics/btx094.

Tada T, Obata Y, Tada M, Goto Y, Nakatsuji N, Tan S, Kono T, Takagi N. Imprint switching for non-random X-chromosome inactivation during mouse oocyte growth. Development. 2000 Jul; 127(14):3101–3105. https://journals.biologists.com/dev/article/127/14/3101/40816/Imprint-switching-for-non-random-X-chromosome, doi: 10.1242/dev.127.14.3101.

Wagner KU, Wall RJ, St-Onge L, Gruss P, Wynshaw-Boris A, Garrett L, Li M, Furth PA, Hennighausen L. Cre-mediated gene deletion in the mammary gland. Nucleic Acids Research. 1997 Nov; 25(21):4323–4330. https://academic.oup.com/nar/article-lookup/doi/10.1093/nar/25.21.4323, doi: 10.1093/nar/25.21.4323.

Wang CY, Colognori D, Sunwoo H, Wang D, Lee JT. PRC1 collaborates with SMCHD1 to fold the X-chromosome and spread Xist RNA between chromosome compartments. Nature Communications. 2019 Dec; 10(1):2950. http://www.nature.com/articles/s41467-019-10755-3, doi: 10.1038/s41467-019-10755-3.

Wang CY, Jégu T, Chu HP, Oh HJ, Lee JT. SMCHD1 Merges Chromosome Compartments and Assists Formation of Super-Structures on the Inactive X. Cell. 2018 Jul; 174(2):406–421.e25. https://linkinghub.elsevier.com/retrieve/pii/S0092867418305841, doi: 10.1016/j.cell.2018.05.007.

Wanigasuriya I, Gouil Q, Kinkel SA, Tapia del Fierro A, Beck T, Roper EA, Breslin K, Stringer J, Hutt K, Lee HJ, Keniry A, Ritchie ME, Blewitt ME. Smchd1 is a maternal effect gene required for genomic imprinting. eLife. 2020 Nov; 9:e55529. https://elifesciences.org/articles/55529, doi: 10.7554/eLife.55529.

Wutz A, Jaenisch R. A Shift from Reversible to Irreversible X Inactivation Is Triggered during ES Cell Differentiation. Molecular Cell. 2000 Apr; 5(4):695–705. https://linkinghub.elsevier.com/retrieve/pii/S1097276500802488, doi: 10.1016/S1097-2765(00)80248-8.

Xue Y, Zhong Z, Harris CJ, Gallego-Bartolomé J, Wang M, Picard C, Cao X, Hua S, Kwok I, Feng S, Jami-Alahmadi Y, Sha J, Gardiner J, Wohlschlegel J, Jacobsen SE. Arabidopsis MORC proteins function in the efficient establishment of RNA directed DNA methylation. Nature Communications. 2021 Dec; 12(1):4292. http://www.nature.com/articles/s41467-021-24553-3, doi: 10.1038/s41467-021-24553-3.

Zeng TB, Pierce N, Szabó PE. H3K9 methyltransferase EHMT2/G9a controls ERVK-driven non-canonical imprinted genes. bioRxiv. 2021 Mar; http://biorxiv.org/lookup/doi/10.1101/2021.03.29.437617, doi: 10.1101/2021.03.29.437617.

